# Vglut2 expression in dopamine neurons contributes to post-lesional striatal reinnervation

**DOI:** 10.1101/2019.12.23.887323

**Authors:** Willemieke M. Kouwenhoven, Guillaume Fortin, Anna-Maija Penttinen, Clélia Florence, Benoît Delignat-Lavaud, Marie-Josée Bourque, Thorsten Trimbuch, Milagros Pereira Luppi, Jean-François Poulin, Christian Rosenmund, Raj Awatramani, Louis-Éric Trudeau

## Abstract

In Parkinson’s disease, the most vulnerable neurons are found in the ventral tier of the substantia nigra (SN), while the adjacent dopamine (DA) neurons of the ventral tegmental area (VTA) are mostly spared. Although a significant subset of adult VTA DA neurons expresses *Vglut2*, a vesicular glutamate transporter, and release glutamate as a second neurotransmitter in the striatum, only very few adult SN DA neurons have this capacity. Previous work has demonstrated that lesions created by neurotoxins such as MPTP and 6-hydroxydopamine (6-OHDA) can upregulate the expression of Vglut2 in surviving DA neurons. Currently, the molecular mechanisms explaining the plasticity of Vglut2 expression in DA neurons are unknown, as are the physiological consequences for DA neuron function and survival. Here we aimed to characterize the developmental expression pattern of Vglut2 in DA neurons and the role of this transporter in post-lesional plasticity in these neurons. Using an intersectional genetic lineage-mapping approach, based on Vglut2-Cre and TH-Flpo drivers, we first found that more than 98% of DA neurons expressed *Vglut2* at some point in their embryonic development. Expression of this transporter was detectable in most DA neurons until E11.5 and was found to be localized in developing axons. Moderate enhancement of VGLUT2 expression in primary DA neurons caused an increase in axonal arborization length. Compatible with a developmental role, constitutive deletion of Vglut2 caused a regional defect in TH-innervation of the dorsal striatum in E18.5 embryos. Moreover, using an *in vitro* neurotoxin model, we demonstrate that *Vglut2* expression can be upregulated in post-lesional DA neurons by 2.5-fold, arguing that the developmental expression of *Vglut2* in DA neurons can be reactivated at postnatal stages and contribute to post-lesional plasticity of dopaminergic axons. In support of this hypothesis, we find fewer mesostriatial dopaminergic projections in the striatum of conditional Vglut2 KO mice 7 weeks after a neurotoxic lesion, compared to control animals. Thus, we propose here that one of the functions of Vglut2 in adult DA neurons is to promote post-lesional recovery of meso-striatal axons.

## INTRODUCTION

In Parkinson’s disease, the most vulnerable neurons are found in the ventral tier of the substantia nigra (SN), while the adjacent dopamine (DA) neurons of the ventral tegmental area (VTA) are mostly spared. Although a significant subset of adult VTA DA neurons expresses *Vglut2*, a vesicular glutamate transporter, and release glutamate as a second neurotransmitter in the striatum, only very few adult SN DA neurons have this capacity (Bérubé-Carrière et al., 2012; Mendez et al., 2008; Poulin et al., 2018; Yamaguchi et al., 2013, 2015). Adult VTA DA neurons that co-express *Vglut2* project to the medial shell of the nucleus accumbens, while the few adult SN DA-Vglut2 neurons project densely to the tail of the striatum and more sparsely in rostral regions (Poulin et al., 2018). Interestingly, the level of *Vglut2* transcript in adult DA neurons is very low i.e. 10-15 copies per cell, which is approximately ten-fold lower than the number of copies of *Th* mRNA or about 5-fold lower than *Vmat2* transcript, in the same cells (Li et al., 2013). Moreover, while this study did not compare the levels of *Vglut2* expression to the levels present in pure glutamatergic neurons of other *bona fide* glutamatergic structures, other qualitative *in situ* hybridization experiments revealed a much higher expression of *Vglut2* in these pure glutamatergic neurons (Yamaguchi et al., 2007). Thus, while these VTA DA neurons co-express *Vglut2*, they still display a marked preference for their dopaminergic identity. The exact role of *Vglut2* expression, and why it is expressed at this level, is currently unclear, although several gain- and loss-of-function papers suggested that the presence of Vglut2 could contribute to 1) axonal outgrowth (Fortin et al., 2012; Schmitz et al., 2009), and 2) striatal DA and (co-) glutamate release (Fortin et al., 2012; Hnasko et al., 2010), and 3) cell survival (Dal Bo et al., 2008; Shen et al., 2018; Steinkellner et al., 2018). In line with this first possibility, cultured DA neurons from Vglut2 conditional knock-out (Vglut2cKO) mice display a smaller axonal arborization (Fortin et al., 2012). Moreover, the NMDA receptor subunit NR1 is present on the axon growth cone of cultured DA neurons, and DA neurons grown in the presence of a NR1 agonist display enhanced axonal growth (Schmitz et al., 2009). The authors suggested that an autocrine glutamate feedback loop may contribute to this axonal outgrowth.

Compatible with the second of these functions, Vglut2 in DA neurons appears to contribute to a facilitation of DA release in ventral striatal regions. Decreased levels of evoked DA release in the nucleus accumbens were detected by fast scanning cyclic voltammetry in conditional Vglut2 knock-out mice (Fortin et al., 2012; Hnasko et al., 2010). Moreover, it has been proposed that the presence of VGLUT2 on a synaptic vesicle could promote DA synaptic vesicle loading through altering the vesicular pH (Aguilar et al., 2017). It needs to be noted however, that several studies have shown that VGLUT2 is typically found to be segregated from TH, DAT and VMAT2 in dopaminergic axonal varicosities (Fortin et al., 2019; Zhang et al., 2015), minimizing the potential significance of this cellular function of VGLUT2 in adult DA axons.

Finally, Vglut2 is thought to contribute to DA neuron survival, as DA neurons without the presence of Vglut2 are more vulnerable to the neurotoxins 6-hydroxydopamine (6-OHDA) or MPTP (Shen et al., 2018; Steinkellner et al., 2018). Furthermore, 2-weeks after either a 6-OHDA or MPTP lesion, more surviving DA neurons express *Vglut2* (Dal Bo et al., 2008; Shen et al., 2018; Steinkellner et al., 2018). The hypothesis that Vglut2 contributes to survival of DA neurons has in fact been tested by viral over-expression of Vglut2 in DA neurons in an *in vivo* mouse model. While one group reported enhanced survival of DA neurons in MPTP-model following VGLUT2 overexpression (Shen et al., 2018), the other reported a massive cell loss induced due to VGLUT2 overexpression (in absence of a neurotoxin (Steinkellner et al., 2018)).

In the current study, we aimed to clarify the cellular roles of Vglut2 in DA neurons. We show that *Vglut2* is expressed in nearly all DA neurons during embryonic development up to the developmental period during which their axons reach target cells in the striatum (Kolk et al., 2009), and that moderate over-expression of VGLUT2 in cultured DA neurons increases axon size. Furthermore, we provide novel evidence showing that *in vitro* MPP+ treatment enhances *Vglut2* expression by DA neurons and propose that the enhanced *Vglut2*, although not required for the basal survival of DA neurons, contributes to post-lesional reinnervation of the striatum by these neurons.

## RESULTS

### Nearly all DA neurons have a Vglut2-lineage and Vglut2 is expressed at E11.5 in DA neurons

In the adult brain a significant subset of VTA DA neurons expresses *Vglut2*, while only very few adult SN DA neurons express *Vglut2* (Bérubé-Carrière et al., 2012; Mendez et al., 2008; Poulin et al., 2018; Yamaguchi et al., 2013, 2015). In contrast, earlier observations reported that *Vglut2* transcript during late embryogenesis and early neonatal development was more broadly expressed in DA neurons (Dal Bo et al., 2008; Mendez et al., 2008). In order to delineate exactly which DA neurons have a Vglut2-positive lineage, we took advantage of a intersectional genetic approach (Poulin et al., 2018), visualising only DA neurons that have *Vglut2* expression history at P1. Using Vglut2Cre and Th-Flpo as drivers, we visualized only cells with a history of co-expression of *Th* and *Vglut2* by the expression of tdTomato (Fig 1A). Using TH protein as a marker for DA neurons, we determined that almost all SN (99%, ± 0.3, n=3) and VTA DA neurons (98%, ±0.3, n=3) have a Vglut2-lineage (Fig 1B). A similar intersectional approach using Dat-TTA and Vglut2-Cre as conditional drivers equally resulted in high percentages of Vglut2-lineage in DA neurons, although slightly lower than with the Th-driver (Fig S1). This is most likely explained by known lower levels of Dat expression in VTA DA neurons (Poulin et al., 2014; Veenvliet et al., 2013), which could lead to underestimation of the number of DA neurons with a Vglut2-lineage. Interestingly in these experiments, we also noted the presence of a subpopulation of neurons with Vglut2-lineage that were TH-negative, suggesting that the dopaminergic phenotype in these neurons was down-regulated below detection level (Fig 1C, white arrow). Together, these experiments indicate that practically all DA neurons have expressed *Vglut2* before P1. These data complement and extend a recent study showing, using a general Vglut2-Cre reporter mouse, that most adult DA neurons possess a Vglut2-lineage (Steinkellner et al., 2018).

**Figure 1:**
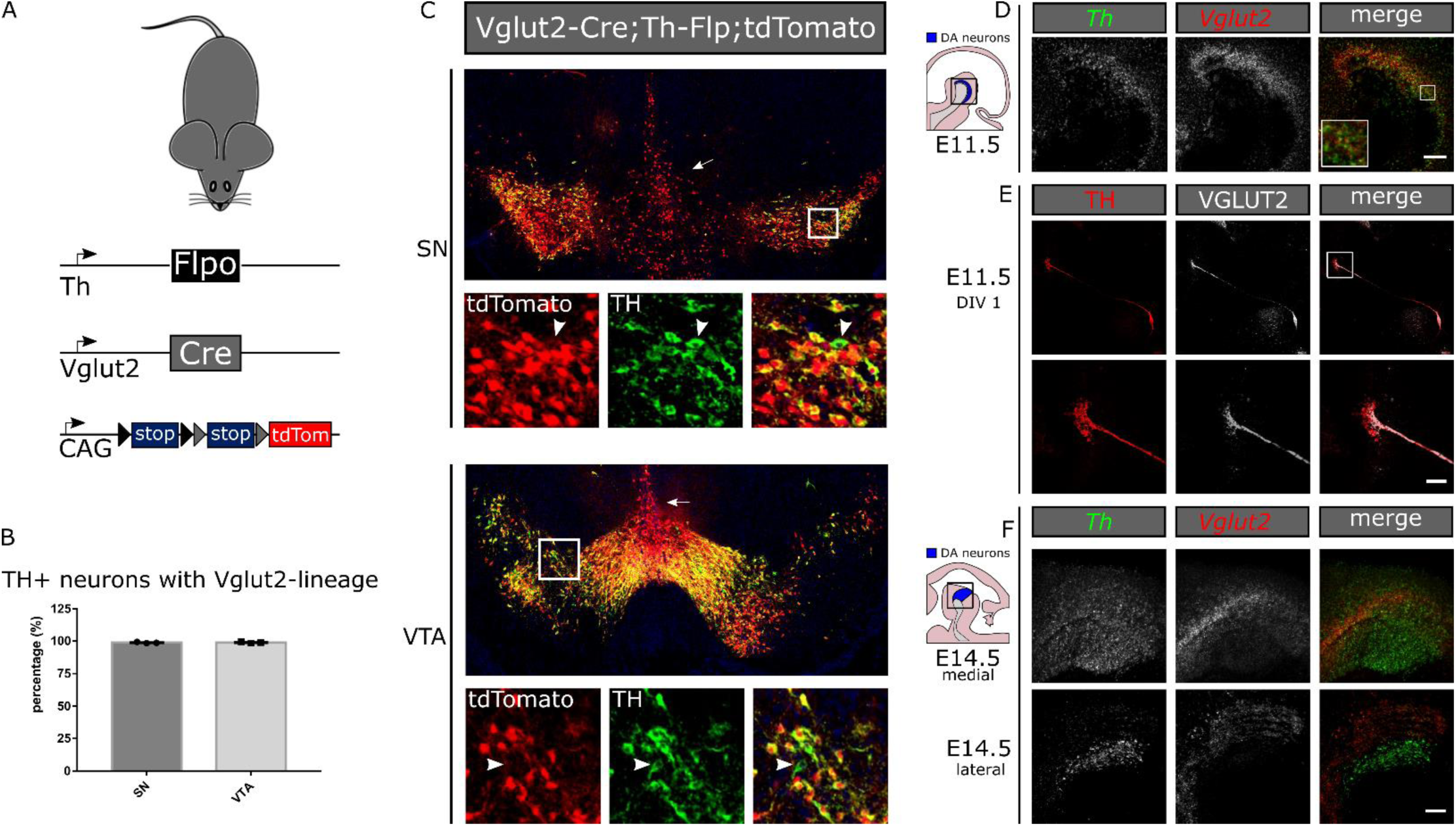
Most DA neurons have a Vglut2 expression history. A. Schematic depiction of the intersectional genetic approach used: mice expressing a Th-Flpo and Vglut2-Cre conditional construct will express tdTomato only in cells that have expressed both Th and Vglut2 genes. B-C. 98% of SN and VTA TH-positive neurons co-localize with tdTomato. Only a small number of TH-positive neurons are negative for tdTomato (white arrowhead). In the medial midbrain, tdTomato positive neurons are found, that are no longer positive for TH (white arrow). D. Fluorescent *in situ* hybridization reveals that Vglut2 expression overlaps with that of Th in the mesencephalon at E11.5. Scale bar represents 100µm E. THGFP-positive neurons taken at E11.5 from the mesencephalon express abundant Vglut2 protein after 24h in culture (DIV1), both in the soma and growth cone (white box). Scale bar represents 10µm F. Vglut2 transcript is still present in the mesencephalon at E14.5, and only partly overlaps with Th transcript in medial sections of the mesencephalon, but not in lateral sections. Scale bar represents 25µm.

In order to determine at what developmental stage *Vglut2* is expressed in DA neurons, we used double fluorescent *in situ* hybridization and observed overlapping *Vglut2* and *Th* expression patterns at E11.5 in the mesencephalon (Fig 1D). Moreover, to establish if VGLUT2 protein is also found in DA neurons at this age, we dissected the mesencephalon of E11.5 pups from animals that express GFP under the TH-promoter, and selecting TH-positive cells using fluorescence activation cell sorting (FACS). Consistent with the *Vglut2* transcript data, we observed clear VGLUT2 protein THGFP-positive DA neurons that were grown *in vitro* for 1 day (DIV1) (Fig 1E). VGLUT2 protein was located both in TH-positive cell bodies, as well as in the growth cones of these developing neurons (Fig 1E, insert).

Interestingly, at E14.5 the transcript expression pattern of *Vglut2* no longer overlapped completely with the expression of *Th*. *Vglut2* transcript was still present in *Th*-positive cells located in medial sections of the mesencephalon but was no longer present in lateral sections of the mesencephalon (Fig 1F). Our findings are also compatible with a recent report describing a higher degree of overlapping expression of *Vglut2* and *Th* at E11.5 in the mesencephalic flexure, compared to E14.5 (Dumas and Wallen-McKenzie, 2019).

### Vglut2 promotes axonal growth in cultured DA neurons

Early expression of Vglut2 in the growth cone of DA neurons during embryonic development is compatible with a potential role of glutamate release in the axonal development and connectivity of DA neurons. At E11.5 DA neurons start extending their axons dorsally and rostrally, and at E14.5 the first growth cones reach the striatum (Kolk et al., 2009). We recently showed that dorsal striatal target cells exert a repressive effect on Vglut2 expression in DA neurons (Fortin et al., 2019). In order to test the role of VGLUT2 expression in DA neurons on axon outgrowth, we investigated the effect of moderate over-expression of VGLUT2, using a lentiviral Vglut2-Venus construct driven by a synapsin promoter, in a monoculture of primary postnatal DA neurons (Fig 2A). Quantification of VGLUT2 protein immunoreactivity in these primary DA neurons by confocal microscopy captured images of primary DA neurons confirmed that this approach increased the levels of VGLUT2 protein by approximately 50% (Fig 2B), thus staying within a physiological range of expression. Next, using confocal microscopy and MAP2 immunostaining to distinguish between axons (MAP2-negative) and dendrites (MAP2-positive), we reconstructed the complete axonal arborization of the DA neurons (Fig 2C, 2D). DA neurons over-expressing Vglut2-Venus displayed a significantly larger axonal arbor compared to control DA neurons transfected with empty Venus-vector (6672 ± 739.4 pixels, n=37 vs 9423 ± 1006 pixels, n=40, Fig 2E). Moreover, the axonal arborization of DA neurons expressing Vglut2-Venus was more complex, based on the increased number of branches (Fig 2F), as well as higher scores in a Scholl analysis (Fig S3). Interestingly, dendritic size, shape and complexity were not affected by the over-expression of Vglut2-Venus (Fig 2). These data are in line with previous observations from our group, showing that cultured DA neurons display smaller axonal arborizations in absence of Vglut2 (Fortin et al., 2012). To test if the over-expression of Vglut2 had effects on DA neuronal survival, we treated infected cultures with 1 µM MPP+ and quantified the number of surviving cells. We observed no differences in survival between DA neurons over-expressing Vglut2-Venus compared to Venus expressing DA neurons, neither in the control condition nor in the MPP+ treated condition (Fig S3). Thus, we conclude that one of the early roles of Vglut2 expression in cultured DA neurons may be to contribute to the development and connectivity of their axonal arbors, and that Vglut2 is not directly involved in DA neuron cell survival at this early time point.

**Figure 2:**
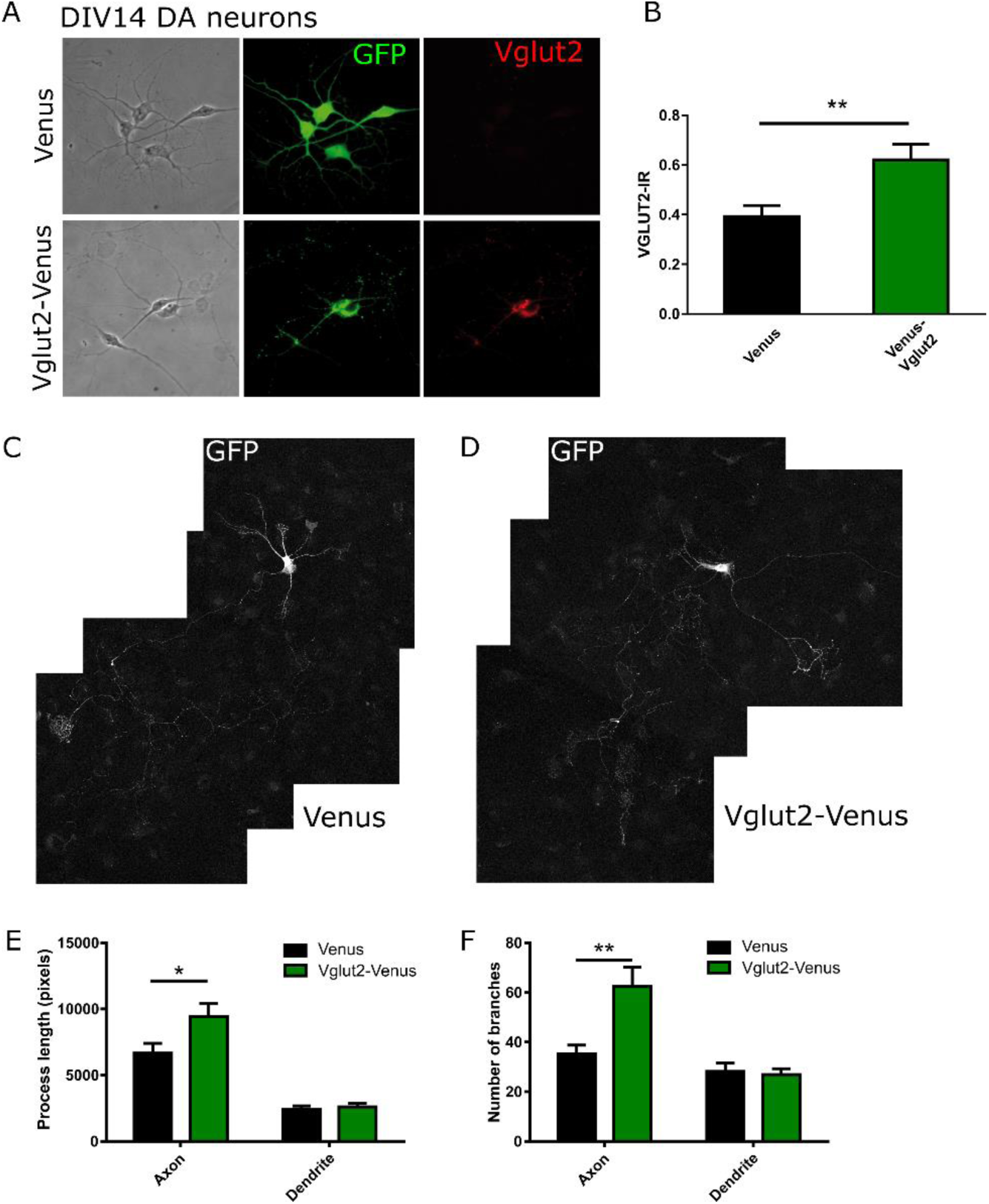
Vglut2 promotes axonal growth in DA neurons *in vitro*. A. THGFP-positive mesencephalic neurons were cultured at P1 for 14 days (DIV14) and infected with Vglut2-Venus lentivirus or control Venus lentivirus. Vglut2 protein is detectable in Vglut2-Venus infected neurons. B. Quantification of Vglut2 protein levels revealed a 50% increase in immunoreactivity (n=53-54, P<0.01, Student’s t-test). C-F. Mesencephalic neurons over-expressing Vglut2-Venus display a larger axonal arborization and increased number of branches. C-D GFP protein expression was used to visualize the general morphology of the cells. E. Quantification of the axonal arbor reveals an enhanced arbor in neurons overexpressing Vglut2-Venus (n=37-40 neurons, P<0.05, two-way ANOVA). F. Branching analysis shows enhanced complexity in axonal arborization due to Vglut2-Venus overexpression (n=37-40 neurons, P<0.01, two-way ANOVA). *= P<0.05, **= P<0.01

### Regional striatal innervation deficit in absence of Vglut2

Knowing that Vglut2 is expressed in the vast majority of DA neurons at E11.5, and that Vglut2 contributes to axon outgrowth in these cells (Fig 1-2), we re-examined the anatomy of the DA system in absence of Vglut2. Previous loss-of-functions studies removed Vglut2 conditionally in DA neurons using the DA transporter (Dat/Slc6a3) as a driver (Fortin et al., 2012; Hnasko et al., 2010). Dat is expressed starting E14, however Cre recombinase activity was observed starting E15 in similar genetic approaches (Bäckman et al., 2006). Consequently, using this approach, Vglut2 expression is deleted after the major peak of Vglut2 expression in DA neurons. For this reason, we investigated the DA system in constitutive Vglut2 knock-out (Vglut2KO mice) at E18.5, as these animals display perinatal mortality (Wallén-Mackenzie et al., 2006). Using TH immunohistochemistry, we observed no overt changes in the macroscopic organization of the mesotelencephalic DA system in Vglut2KO mice compared to Vglut2WT littermate controls (Fig 3A). Furthermore, quantification of TH-positive neurons revealed no difference in the number of DA neurons in the mesencephalon of Vglut2KO E18.5 embryos (Fig. 3B), demonstrating that embryonic Vglut2 expression in DA neurons is not directly required for DA neuronal cell survival.

**Figure 3:**
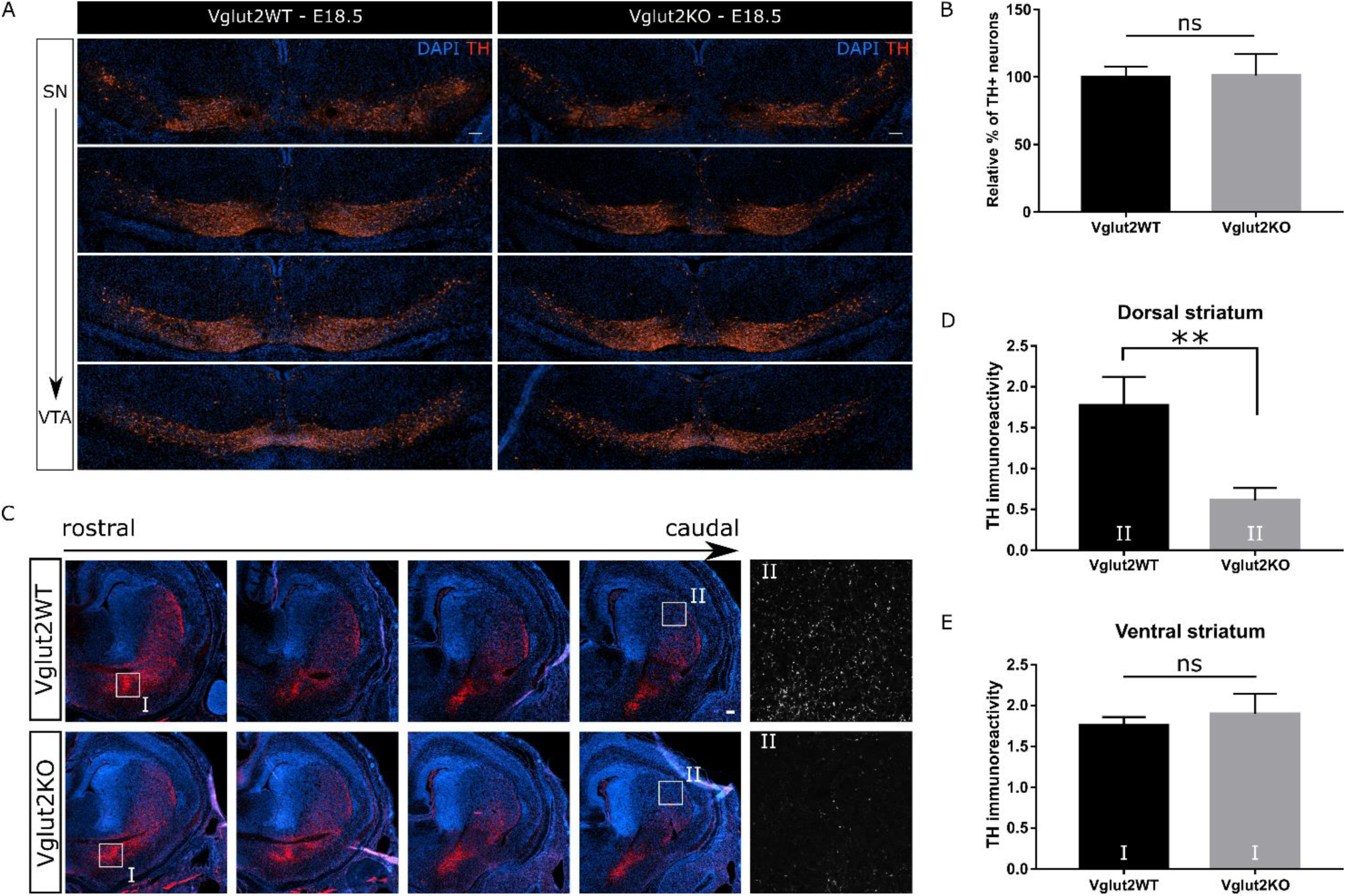
Unaltered DA neuronal development in absence of Vglut2, but regional striatal innervation deficit. A. immunohistochemistry analysis against TH and DAPI in coronal slices of E18.5 Vglut2WT and constitutive Vglut2KO littermates reveals no macroscopic changes in the dopaminergic system in the mesencephalon. Scale is 100 µm. B. Quantification of the number of TH-positive neurons at E18.5 in the mesencephalon reveals no significant differences between Vglut2WT and Vglut2KO littermates. C. immunohistochemistry analysis against TH and DAPI in coronal slices of E18.5 Vglut2WT and Vglut2KO littermates reveals no macroscopic changes in the striatum. Scale is 100 µm. D-E. Quantification of TH-immunoreactivity in the striatum at E18.5, reveals a regional decrease of in intensity in caudal, dorsal striatum (D: n=4 pups, P<0.01, Student’s t-test) but not in ventral striatum (E: n=4 pups, P>0.05, Student’s t-test). **= P<0.01

As we and others observed a role for Vglut2 and glutamate in axonal growth (Fortin et al., 2012; Schmitz et al., 2009), we investigated whether the striatal innervation of DA neurons was altered in absence of Vglut2 by quantifying the density of TH immunoreactivity in different sectors of the striatum. We observed no overt macroscopic changes in the development of DA striatal innervation in Vglut2KO compared to Vglut2WT littermate controls (Fig 3C). However, in the primordial regions that correspond to the adult target regions of Vglut2-expressing DA neurons in the SN and VTA, i.e. the caudal, dorsal striatum and the ventral striatum, respectively (Poulin et al., 2018), we observed a significant decrease in TH-immunoreactivity in the caudal dorsal striatum, but not in the ventral striatum (Fig 3C-D). This suggests that although in general the striatum is successfully innervated by DA projections in the absence of Vglut2, some of the projections are hampered in their development.

### Th and Vglut2 expression in DA neurons are highly plastic and regulated in opposite ways

The gradual decrease of *Vglut2* expression by DA neurons during embryonic development suggests that intrinsic or extrinsic signals regulate expression of this gene in DA neurons. We previously showed that interaction of DA neurons with dorsal striatal cells *in vitro* appears to negatively regulate *Vglut2* expression in these neurons (Fortin et al., 2019). Other work has shown that neurotoxic lesions increase the likelihood of *Vglut2* expression in surviving DA neurons (Dal Bo et al., 2008; Shen et al., 2018; Steinkellner et al., 2018), but it is currently not yet clear whether this is caused by enhanced *Vglut2* expression per DA neuron or enhanced survival of DA neurons that co-express *Vglut2* and *Th*. As cellular stress frequently reactivates previously suppressed developmental programs, here, we examined the possibility that MPP+, a toxin often used to mimic Parkinson’s disease-related cellular stress, might trigger increased expression of *Vglut2* in parallel to its well-known ability to reduce *Th* expression (Bowenkamp et al., 1996). To test this hypothesis we treated cultured SN neurons, taken from DatCre;AI9 neonatal pups (that express tdTomato in Dat-positive neurons) and treated the cells with a low dose (2µM) of MPP+. After 72h, we FAC-sorted the tdTomato-positive DA neurons (Fig 4A). Quantification of the number of FACS-purified neurons, demonstrated no significant cell loss after the 2µM MPP+ treatment, confirming the chosen dose to be sub-lethal (n = 3 cultures, Student’s t-test, P>0.05, Fig 4B).

**Figure 4:**
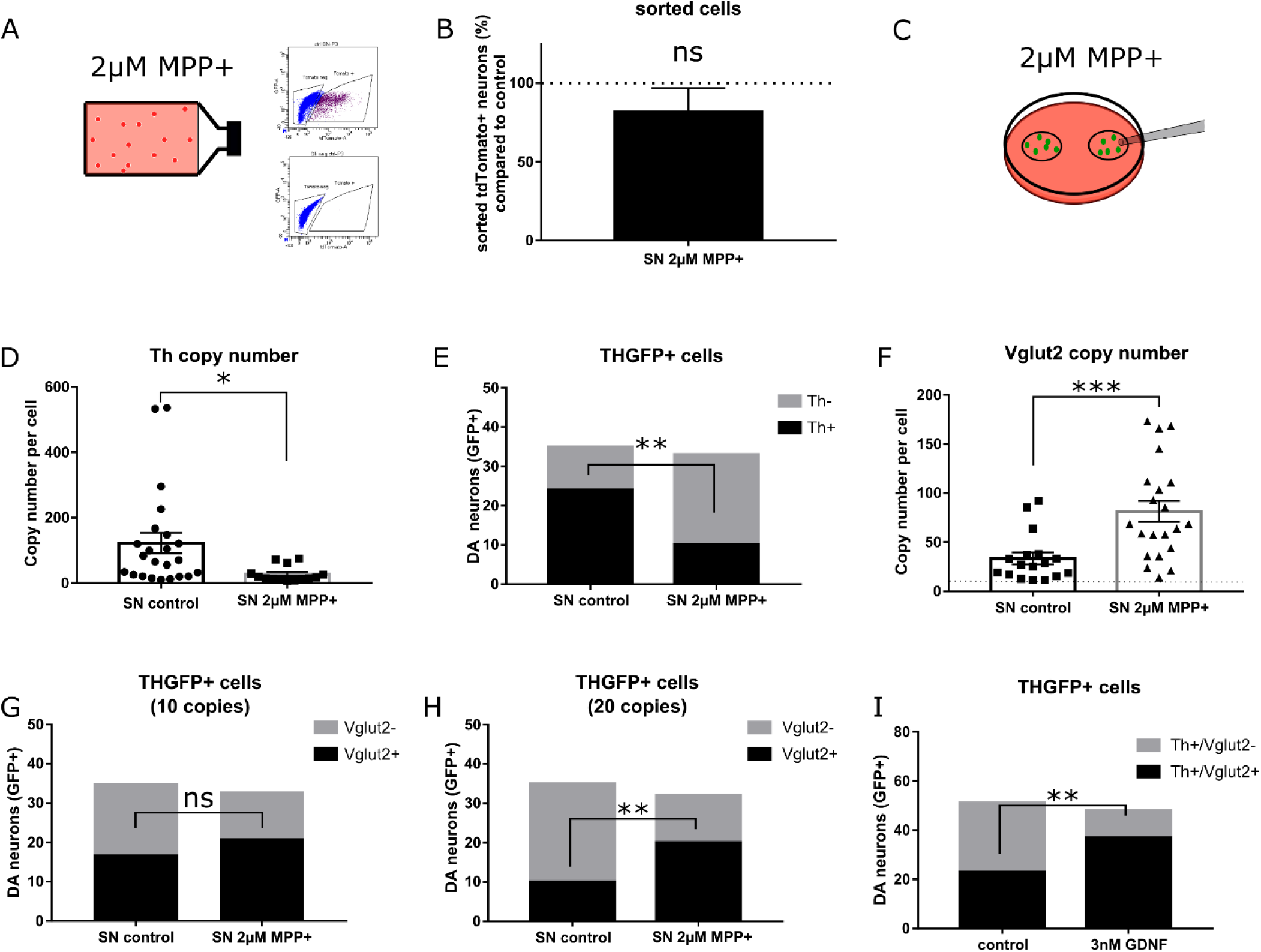
Th and Vglut2 expression in DA neurons are highly plastic and are regulated in opposite ways. A. THGPP-positive SN neurons were grown in a petri-dish and were collected with a glass pipet at DIV14, after 72h of 2µM MPP+ treatment. B. SN neurons expressed on average 122 (± 31, n=35) copies of *Th* transcript at DIV14, and this decreased to 30 (± 7, n=12) copies after MPP+ treatment (Student’s t-test, P<0.05). F. Significantly fewer SN neurons expressed detectable levels of *Th* mRNA after MPP+ treatment (12/33, Fisher’s exact test), compared to control (23/35, Fisher’s exact test P<0.01). SN neurons contained on average 33 (± 6 copies, n=35) of *Vglut2* transcript in at DIV14, and this increased to 81 (± 11, n=33) copies after MPP+ treatment. G. No changes were observed in the number of SN neurons that express *Vglut2* in control (17/35, Fisher’s exact test, n.s.) compared to MPP+ treatment (21/33). H. Cultured DA neurons that were treated for 3h with GDNF (3nM) before being collected were significantly more likely to contain both Vglut2 and Th transcripts compared to DA neurons in untreated cultured (n=40 cells, 4 cultures, Fisher’s exact test, P<0.01). *= P<0.05, **= P<0.01

Next, in order to determine the exact copy number of *Th* and *Vglut2* present per SN neuron, both in control and after a sub-lethal dose of MPP+, we repeated this paradigm, but collected individual GFP-positive DA neurons, taken from a primary neuronal culture prepared from dissected SN neurons of THGFP-positive neonatal pups. Again, at DIV11 we treated the cells with 2µM of MPP+ for 72h, before we collected the cells using a glass pipet, after which we performed single-cell qPCR (Fig 4C). We found that MPP+ caused a significant decrease in *Th* transcript expression per SN neuron (Fig 4D, 122 ± 31 copies, n=23 vs 30.44 ± 7 copies, n=12 cells, Student’s t-test, P<0.05). Moreover, the proportion of THGFP-positive neurons that still expressed *Th* above threshold detection level (i.e. 10 mRNA copies) decreased significantly after the MPP+ treatment (24/35 neurons (69%) vs 10/34 neurons (30%), P<0.01) (Fig. 4E). As previously mentioned, *Vglut2* transcript levels in adult DA neurons *in vivo* are quite low, i.e. 10 mRNA copies per DA neuron (Li et al., 2013). Using a similar inclusion criterion of 10 copies of *Vglut2* mRNA to classify a DA neuron as Vglut2+, we first observed that GFP-positive SN neurons displayed a more than 2-fold increase in *Vglut2* mRNA copy number after MPP+ treatment (Fig 4F, 33 ± 6, n=17, 81 ± 11, n=21, P<0.001). We next examined if the treatment also changed the proportion of SN DA neurons containing *Vglut2* mRNA. In control, untreated neurons, almost 50% of the control SN neurons were Vglut2+ positive (17/35 neurons) and contained more than 10 copies of *Vglut2* mRNA. After MPP+ the percentage of DA neurons containing more than 10 copies did not change (17/35 neurons (49%) vs 21/33 neurons (64%), n.s.) (Fig. 4H). However, it is striking to note that with a more stringent criterion of 20 mRNA copies, the MPP+ treatment caused a large increase in the proportion of Vglut2+ DA neurons (10/35 neurons (28%) for controls vs 21/33 neurons (62%) after MPP+ treatment, P<0.01) (Fig 4G). Together these data suggest that a large subset of SN DA neurons have the potential to enhance *Vglut2* expression upon exposure to cellular stress induced by MPP+.

Although an enhanced occurrence of *Vglut2* in surviving DA neurons after partial lesions has previously been reported (Dal Bo et al., 2008; Shen et al., 2018; Steinkellner et al., 2018), the specific signals that regulate *Vglut2* expression in DA neurons in response to cellular stress are presently unknown and could be different from developmental regulation of *Vglut2* expression. A striking change that has been observed in the post-lesional brain, is the enhanced expression of growth factor GDNF (Hidalgo-Figueroa et al., 2012). GDNF is normally known to be expressed by parvalbumin-positive striatal interneurons, which are responsible for 95% of the striatal GDNF production. Moreover, GDNF enhances glutamate release in cultured mesencephalic DA neurons (Bourque and Trudeau, 2000). In order to test if GDNF could drive enhanced *Vglut2* expression in DA neurons, we treated cultured DA neurons for 3h at DIV7 with 3nM GDNF. In accordance with this hypothesis, using RT-PCR, we found that the proportion of DA neurons containing *Vglut2* mRNA increased after a 3h treatment with 3nM GDNF, increasing from 45% of the THGFP neurons in control conditions to 77% after GDNF (Fig 4I, Fisher’s exact test; P<0.01).

To recapitulate, we find here that expression of *Th* and *Vglut2* is reciprocally regulated in DA neurons in response to cellular stress. While previous work showed that the presence of striatal cells reinforces the *Th* phenotype of DA neurons (Fortin et al., 2019), a low dose of MPP+ results in the opposite and favors their glutamatergic phenotype, as revealed by an increase in *Vglut2* mRNA copy number. Finally, our *in vitro* data suggest the possibility that upregulated GDNF expression in the post-lesional brain could elevate *Vglut2* expression above detection level and as such could be involved in the observed enhanced prevalence of *Vglut2* expression in DA neurons in the post-lesional brain *in vivo*.

### Fewer striatal-mesencephalic projections after a partial lesion in absence of Vglut2

Based on our results showing the presence of VGLUT2 in the growth cones of DA neurons at early developmental stages, our finding of increased axon development following overexpression of VGLUT2 and our finding of elevated expression of *Vglut2* in DA neurons in response to cellular stress, we next wondered if the increased *Vglut2* expression in DA neurons after a lesion could contribute to post-lesional compensatory axonal outgrowth of these neurons, promoting reinnervation of the striatum. To test this hypothesis we performed an intra-striatal 6-OHDA injection to partially lesion the mesencephalon of adult conditional Vglut2cKO mice and controls (as per a previously published protocol (Giguère et al., 2019)), and examined the innervation pattern of SN and VTA DA neurons after injection of fluorescent retrobeads in the striatum (Fig 5A). We examined the mice 7 weeks after the lesion because reinnervation of the striatum has been observed at this time following a 6-OHDA lesion (Schmitz et al., 2013). We hypothesized that in the absence of Vglut2, DA neurons would be less successful in reinnervation of the striatum than wild-type controls (Fig 5B). To quantify the reinnervation, we injected red fluorescent retrobeads (Rb) at the same coordinates as the 6-OHDA or saline injection (AP 0.5 mm; ML 2.0 mm; DV −3.0 mm), one week before animals were sacrificed. Retrobeads were taken up by terminals and transported to the soma, revealing which midbrain neurons project to the dorsolateral striatum at the 7-week, post-lesion time-point (Fig 5C). Rb-positive neurons were found in both the SN and VTA regions, mainly colocalizing with TH-positive neurons (Fig 5C, yellow arrow), as expected. We nonetheless detected a limited number of labelled neurons which appeared to be TH-negative (Fig 5C, yellow arrowhead). We observed no significant change in the percentage of Rb+/TH-cells (as compared to total Rb+ cells) between conditions (Fig S4A). Using unbiased stereological counting on TH-stained slices, we observed a 48% decrease of the number of SN neurons in controls and a 44% decrease in Vglut2cKO animals (Fig 5D, p<0.05, treatment effect), thus confirming that the injection of 1.5 µg of 6-OHDA (0.5 µL, 3mg/mL) successfully resulted in a partial SN lesion. General analysis of the striatum, using a low-magnification objective, revealed that the stereological cell loss was accompanied by an average decrease of approximately 15% of fluorescent TH-immunoreactivity (TH-IR) in animals of both genotypes that had received a lesion (Fig S4B, p<0.01, F(1,25), n=6-9 animals per condition).

**Figure 5:**
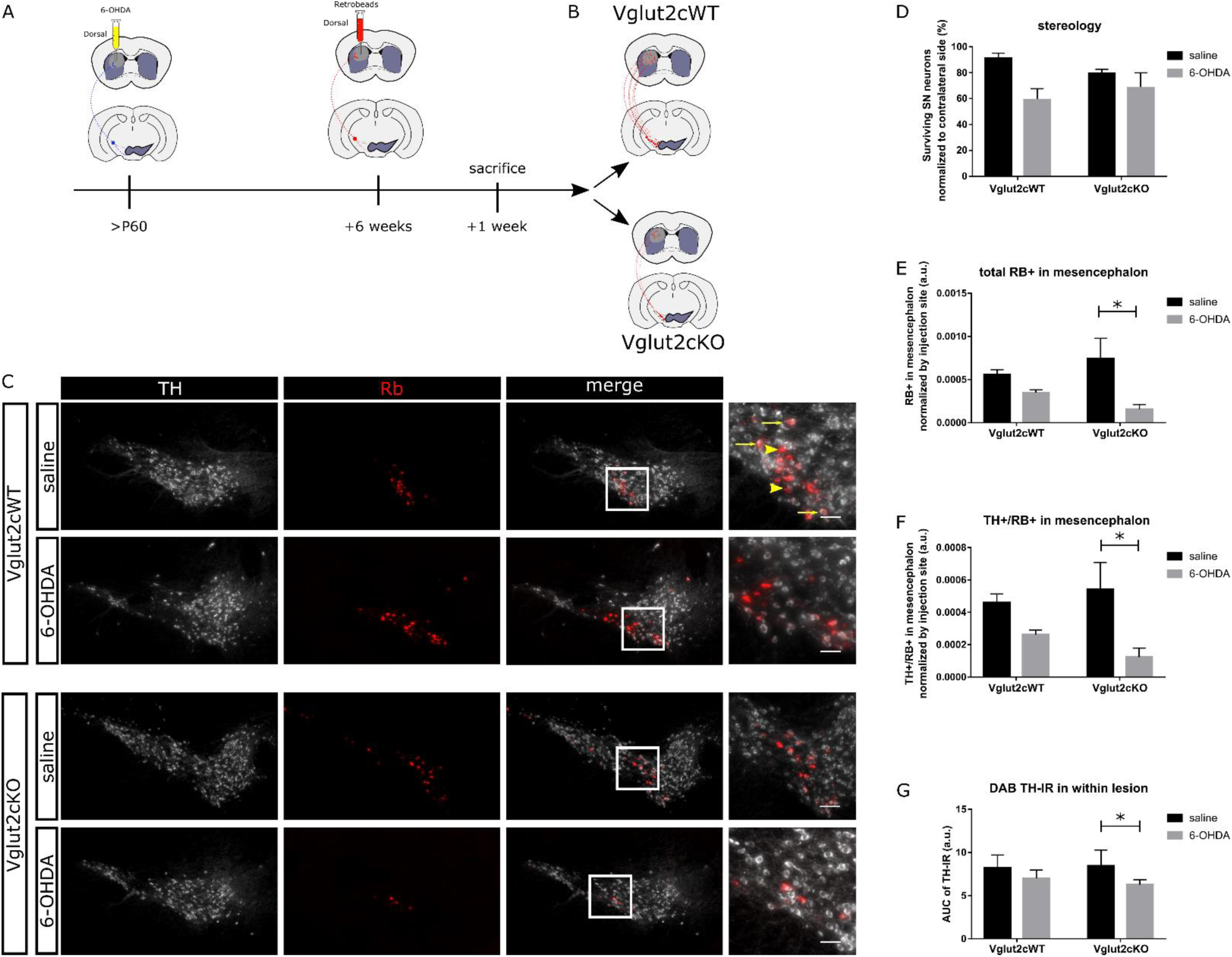
Striatal re-innervation by DA neurons after a partial lesion is perturbed in absence of Vglut2. A. Schematic detailing of the surgery paradigm. B. Schematic representation of the hypothesis that fewer retrobead-positive DA neurons successfully established projections to the dorsal striatum in Vglut2cKO mice after a 6-OHDA lesion compared to wild-type control mice. C. Immunohistochemistry for TH (white signal) reveals the presence of red-fluorescent retrobead-positive cells in the SN, which were typically positive (yellow arrow) or more rarely negative for TH-immunoreactivity (white arrow). Scale is 200µm. D. Unbiased stereological counting on sections stained for TH and cresyl violet confirmed loss of SN TH-IR neurons but revealed no increased sensitivity of Vglut2-ablated SN neurons to 6-OHDA ((treatment effect: F (1, 24) = 6.41, P<0.05, n=5-7 animals per condition) E. Fewer retrobead-positive neurons are observed in mesencephalon of Vglut2cKO animals, after 6-OHDA lesion compared to saline controls (treatment effect: F (1, 20) = 13.99, P<0.01), n=5-8 animals per condition. F. Fewer TH+/RB+ neurons are observed in mesencephalon of Vglut2cKO animals, after 6-OHDA lesion compared to saline controls (treatment effect: F (1, 20) = 14.92, P<0.01), n=5-8 animals per condition. G. Quantification of striatal DAB TH-IR in total lesion per brain confirmed a significant decrease of TH innervation in Vglut2cKO animals after 6-OHDA injections (treatment effect: F (1, 24) =12.17, P<0.01, n=6-9). Tukey’s multiple comparison *P<0.05; **P<0.01.

Next, to assess the success with which striatal-mesencephalic projections had been re-established in the post-lesional brain in the absence of Vglut2, we quantified the total number of Rb-positive neurons as well as Rb/TH-positive neurons in the mesencephalon. To compensate for biased quantification due to variations in injection volume, we divided the number of counted Rb cells by the total surface area of the striatal Rb injection site (Fig S4C). This revealed that in Vglut2cWT control mice, the 6-OHDA lesion caused a modest but non-significant decrease in the number of labelled DA neurons, suggesting the presence of efficient compensatory axonal sprouting (Fig. 5E-F). In contrast, in Vglut2cKO mice the 6-OHDA lesion caused a robust decrease in the number of labelled neurons in mesencephalon (Fig 5E-F), suggesting that fewer striatal-mesencephalic projections were present at this seventh week post-lesion time point in the Vglut2cKO mice. In order to determine whether the absence of Vglut2 equally altered the level of dopaminergic fibers reinnervating the striatum after a lesion, we re-investigated the TH-IR within the lesioned area, with a higher magnification objective, in more detail. Using TH-DAB immunohistochemistry (thus avoiding non-specific fluorescence within the lesioned area caused by the Rb-injection), we quantified the total dorsal striatal lesion size per animal by plotting the intensity of DAB TH-IR per slice and calculating the area under the curve ((AUC), Fig S4D). Compatible with the observation that fewer striatal-mesencephalic projections were present at this seventh week post-lesion time point in the Vglut2cKO mice, we observed a decrease in total DAB TH-immunoreactivity within the lesioned area in both genotypes (F(1,25)=12.43, p<0.01)), but post-hoc testing revealed a significant decrease only in Vglut2cKO mice (p<0.05, 8.4 vs 6.3, (Fig 5G). Thus, the same number of surviving DA neurons re-established fewer TH-positive, striatal connections in absence of Vglut2, suggesting that Vglut2 plays an important role in driving the growth of striatal-mesencephalic projections in the post-lesional brain.

## Discussion

In the present work we report that *Vglut2* is expressed early during the embryonic development of mesencephalic DA neurons. Constitutive KO and overexpression experiments further suggest that one of the roles of this vesicular glutamate transporter in DA neurons may be to promote axonal development. We also explored the possibility that such an early developmental expression of *Vglut2* could be reactivated under conditions of cellular stress, in the subset of DA neurons that maintain a minimum of 10 copies of *Vglut2* transcript (Fig 6A). We propose that the reactivated *Vglut2* expression could contribute to post-lesional axonal sprouting. In line with this hypothesis, we find that neurotoxic stress can increase *Vglut2* mRNA copy number of nigral DA neurons and that in the absence of Vglut2, DA neurons show impaired axonal sprouting in response to a partial lesion.

**Figure 6:**
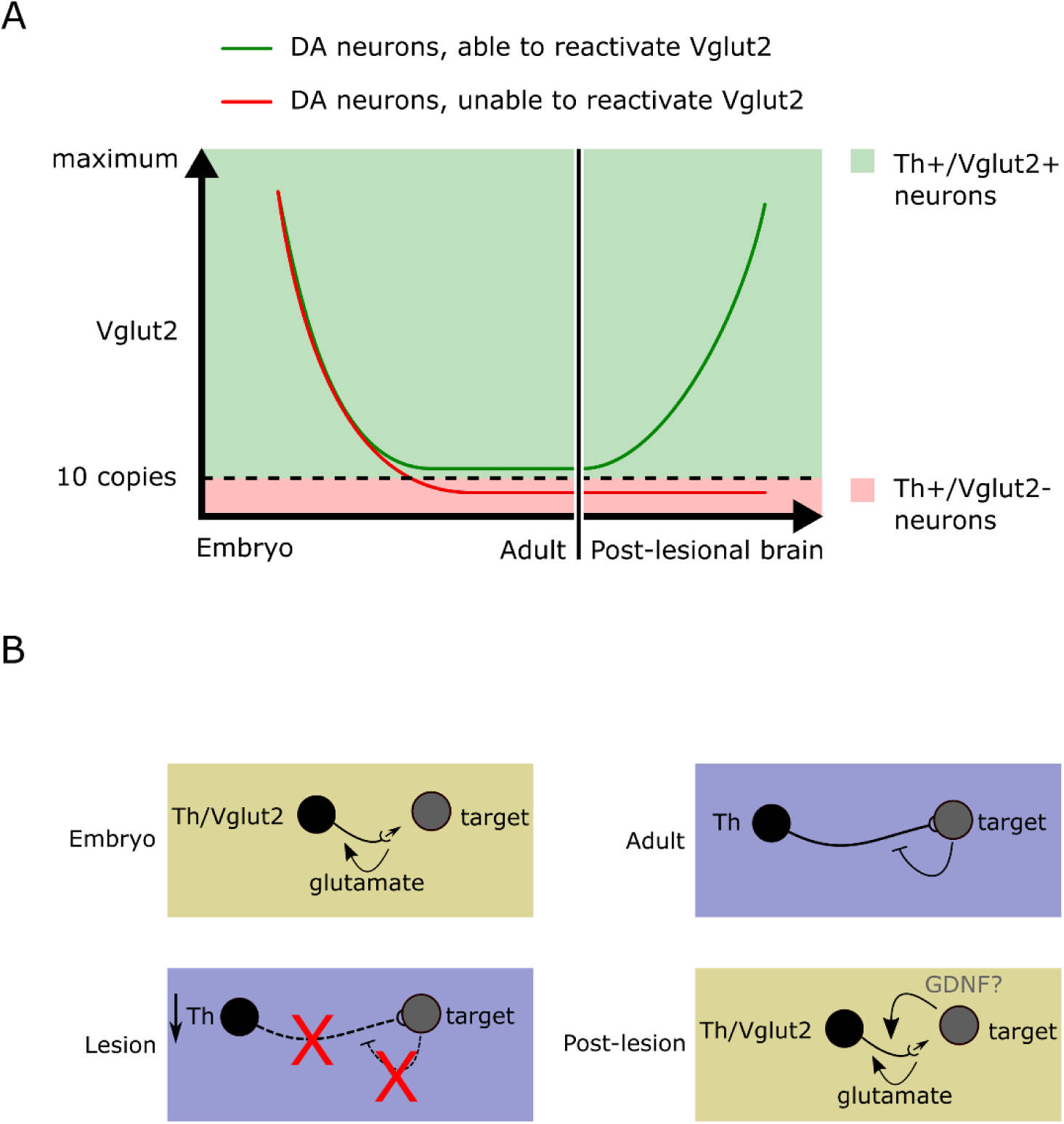
Proposed model of *Vglut2* expression post-lesional plasticity in DA neurons. A. Schematic diagram of a model for the plasticity of Vglut2 expression in DA neurons. Vglut2 is expressed in all DA neurons during early embryonic development, after which it is downregulated during maturation. Based on our single cell qPCR experiments we observed that the subset of DA neurons that maintain Vglut2 expression above the 10 transcript copy threshold are able to reactivate Vglut2 expression after a lesion (red line). In contrast, the subset of DA neurons that downregulate Vglut2 below the 10-copy threshold are not able to reactivate Vglut2 expression in the post-lesional brain (green line). B. Upper Panel. During embryonic development *Vglut2* expression is ubiquitous in DA neurons, allowing for a glutamate autocrine feedback loop, at the time that DA axons are still finding their path towards their striatal target cells. In the adult brain DA neurons have established nigro-striatal connections, at which point the glutamatergic identity of DA neurons is down-regulated. Lower Panel. In response to some major forms of cellular stress, such as neurotoxic lesions, DA axons die back and Th expression is diminished. The loss of nigro-striatal connectivity then lifts this inhibitory feedback on Vglut2 expression in DA neurons. As a result, Vglut2 is upregulated DA neurons in the post-lesioned brain, facilitating the regrowth of axons by DA neurons to regrow their axons and the re-establishment of striatal connections. GDNF expression is increased in the striatum in the post-lesioned brain, and might therefore be involved *in vivo* to enhance vglut2 expression in DA neurons.

### Role and regulation Vglut2 expression in post-lesional plasticity

We previously provided evidence suggesting upregulation of *Vglut2* in surviving DA neurons after a neurotoxic lesion (Dal Bo et al., 2008), a finding recently confirmed by two other groups (Shen et al., 2018; Steinkellner et al., 2018). However, the implications of this observation are presently unclear. Here we first demonstrate that Vglut2 is expressed in essentially all DA neurons during early embryonic development, localizing to the cell body and axonal growth cone, and diminishes shortly after, around E14 (Fig 1). Our findings are in keeping with the results of a micro-array study on Ngn2-GFP sorted DA progenitor cells (Bye et al., 2015), showing high levels of *Vglut2* expression during early embryonic development. Two other recent reports from the same group also confirmed substantial co-expression of *Vglut2* and *Th* at E11.5, although only limited expression of Vglut2 in DA neurons at E14.5 (Papathanou et al., 2018); Dumas and Wallen-McKenzie, 2019). This fluctuation in Vglut2 expression correlates temporally with the moment DA neurons start extending their axons and the moment the first axons of these neurons reach their striatal target cells. Interestingly, it had previously been shown that the ionotropic NMDA glutamate receptor subunit NR1 is found on the growth cone of DA neurons and suggested that axonal outgrowth in primary DA neurons can be driven by an autocrine glutamate-mediated feedback loop (Schmitz et al., 2009). In addition, we have also shown that post-natal dorsal striatal cells repress *Vglut2* expression in cultured DA neurons (Fortin et al., 2019).

In the current work, we examined the plasticity of *Vglut2* expression in primary SN DA neurons using single cell qPCR and we discovered that *Vglut2* levels in single neurons were significantly elevated after a sub-lethal MPP+ treatment. Notably, using an inclusion criterion of 10 mRNA copies, we observed a high prevalence of Vglut2+ SN DA neurons under control conditions. This might reflect the fact that although expression of the *Vglut2* gene is strongly downregulated in postnatal SN DA neurons, a low level of expression persists in a large subset of neurons. In addition, the proportion of Vglut2+ DA neurons may have been partly overestimated, because in order to avoid VTA neurons in the cell collection, we dissected the most lateral and rostral SN neurons (Pacelli et al., 2015), which contains a recently described population of Vglut2+ SN DA neurons (Poulin et al., 2018). Second, the culture was prepared with neonatal DA neurons, which have previously been shown to possess higher levels of *Vglut2* expression compared to more mature DA neurons (Mendez et al., 2008). After the treatment with MPP+, we observed that the proportion of SN DA neurons containing more than 20 copies of *Vglut2* mRNA was significantly increased. Our findings suggest that while it has been a common consensus in the literature that only a small subset of SN DA neurons express Vglut2 in the healthy adult brain (Mendez et al., 2008; Poulin et al., 2018; Yamaguchi et al., 2013), expression of this gene can be re-activated in a larger proportion of DA neurons. Whether all subtypes of SN DA neurons, such as Sox6+, Alhda1a+ or Ndnf+ SN DA neurons (Poulin et al., 2018) are equally likely to re-activate above-threshold expression of *Vglut2* remains to be determined. However, our observation that the proportion of Vglut2+ DA neurons is the same in control and after MPP+ using a 10 mRNA copy number criterion, suggests that only DA neurons that express 10 copies of *Vglut2* or more are able to enhance *Vglut2* expression in response to cellular stress (Fig 6A).

Taken together, these findings lead us to hypothesize that embryonic outgrowth of DA axons is supported by such a glutamate autocrine feedback loop, until mature nigro-striatal connections are established, at which point target cells repress *Vglut2* expression below detection levels, thus inhibiting the autocrine feedback loop (Fig 6B, upper left panel). This repression would be maintained in most DA neurons in adulthood (Fig 6B, upper right panel). In response to some major forms of cellular stress, such as a neurotoxic lesion, we further hypothesize that as *Th* expression is down-regulated in DA neurons and subsets of DA axons die back, the accompanying loss of nigro-striatal connectivity lifts this inhibitory feedback on *Vglut2* expression in DA neurons (Fig 6B, left lower panel). Consequently, *Vglut2* expression could be re-activated in DA neurons and the autocrine loop of glutamate release by DA neurons may be re-instated in axonal growth cones of these neurons. Together, these changes would help the compensatory axonal sprouting program known to promote reinnervation of the partially denervated striatum (Fig 6B, right lower panel). Whether striatal reinnervation is solely driven by Vglut2+ DA neurons, or whether these neurons might function as a guide for Vglut2-DA neurons during the reinnervation process remains to be investigated.

Although speculative at this point, the reactivation of Vglut2 expression in DA neurons could potentially by driven by enhanced levels of neurotrophic factors in the striatum including GDNF. GDNF can promote the double *Vglut2/Th* phenotype in cultured DA neurons (Fig 4) and is augmented in the post-lesional striatum (Hidalgo-Figueroa et al., 2012). Moreover, GDNF is considered dopaminotrophic, as it can boost Th expression, DA release (Hudson et al., 1995) and the formation of axon terminals by DA neurons (Bourque and Trudeau, 2000). In addition, GDNF treatment of ventral midbrain cultures increases neurite outgrowth and complexity (Schaller et al., 2005; Widmer et al., 2000), and over-expression of GDNF in the striatum of non-human primates promotes the neurite outgrowth of embryonic SN grafts (Redmond et al., 2009). These characteristics of GDNF are compatible with the hypothesis that it plays an important role in promoting reinnervation of the striatum by DA neurons in a post-lesional brain. Notably however, *GDNF* transcript has been observed in developing striatum starting at E16.5 (Golden et al., 1999), making it less likely to be also involved in the developmental regulation of *Vglut2* transcript in DA neurons.

While the current study provides support for a role for Vglut2 in axonal outgrowth in DA neurons through axonal glutamate release, it does not exclude that Vglut2 plays other roles in embryonic DA neurons. For example, Vglut2 was first discovered as a Na(+)-dependent inorganic phosphate cotransporter (Aihara et al., 2000; Hisano et al., 2000), and might thus also contribute to cellular homeostasis by regulating phosphate levels. Moreover, the observed VGLUT2 protein expression in cell bodies of embryonic DA neurons (Fig 1) might potentially indicate a role for Vglut2 within the somatodendritic department, as has been proposed for Vglut3 (Fremeau et al., 2004; Kao et al., 2004). Alternatively, enhanced VGLUT2 protein in the cell body at this stage could be a mere side-effect of enhanced Vglut2 transcription and translation, which could equally explain the signal also detected in freshly dissociated adult DA neurons (Mendez et al., 2008).

### Vglut2 expression contributes to axonal outgrowth in DA neurons

In the present work, we studied the role of Vglut2 in post-lesional striatal reinnervation by taking advantage of a partial lesion model targeting nigral neurons and subsequently allowing surviving DA neurons to re-innervate the dorsal striatum. We used a previously published protocol (Giguère et al., 2019), known to create a partial lesion at one month after toxin injection. Our finding of equivalent cell loss in Vglut2cKO mice compared to control mice stands in relative contrast to a recently published report (Steinkellner et al., 2018). An important difference between the lesion protocol used here and the one that is used in the study of Steinkellner and colleagues is the post-lesion time: two months in the present study compared to 10 days in the other report (Steinkellner et al., 2018). It is well-established that the effect of 6-OHDA on DA neurons consists of both down-regulation of TH expression and secondary loss of DA neurons. Several studies demonstrated that the time immediately after a lesion is characterized by down-regulation of TH expression, a cellular phenotype that can be restored, while genuine cell loss is more apparent at later timepoints, (Bowenkamp et al., 1996; Stanic et al., 2003). From that perspective, it could be speculated that DA neurons lacking Vglut2 may down-regulate TH more readily in response to cellular stress compared to control animals, without progressing to cell death.

In the current study, we show that cultured DA neurons overexpressing moderate levels of VGLUT2 grow larger axonal arbors (Fig 1). This observation is in line with our previous loss-of-function study, showing that in the absence of Vglut2, cultured DA neurons grow smaller axons (Fortin et al., 2012). The effect size on axon length is approximately 15%-30% in both the over-expression and the loss-of-function study, highlighting that although playing a regulatory role, Vglut2 is not required for intrinsic axonal outgrowth. Indeed, a large body of literature has shown previously that developmental axonal outgrowth and axon pathfinding of DA neurons depend on an intricate interplay of receptors, ligands and soluble factors (review by (Brignani and Pasterkamp, 2017). Moreover, the potency of the axon guidance factors could also explain why only a modest, regional innervation defect was observed in the constitutive Vglut2KO mice. We thus hypothesize that while DA neurons grow smaller axons in absence of Vglut2, in the otherwise intact embryonic brain, other axon guidance factors still enable correct striatal innervation by most DA neurons. Many of these guidance factors are expected to be absent in our *in vitro* paradigm, and several have been shown to be actually decreased in the post-lesioned brain (Kalaani et al., 2016), thus potentially explaining our observation of a phenotype in cultured Vglut2cKO DA neurons (Fortin et al., 2019), and of the perturbed striatal re-innervation in absence of Vglut2 (Fig 5).

Our observation of increased axonal growth after lentiviral overexpression of VGLUT2, although compatible with previous work showing that glutamate release by the growth cone of DA neurons may promote axonal growth (Schmitz et al., 2009), stands in stark contrast with a recent study showing that overexpressing VGLUT2 in DA neurons leads to massive degeneration of SN DA neurons (Steinkellner et al., 2018). A number of factors need to be considered to explain such different conclusions. First, the culture system used here only investigates the cell intrinsic effects of VGLUT2 overexpression in young, early postnatal DA neurons, such as the previously stated autocrine loop, but does not take into account possible more indirect mechanisms that could influence the survival of adult DA neurons. Second, *Vglut2* transcript levels in adult DA neurons *in vivo* are very low, i.e. 10 *Vglut2* copies per DA neuron (Li et al., 2013). Here, we show that *in vitro* MPP^+^ treatment induced an approximate 2-fold increase of *Vglut2* expression in DA neurons (Fig 4), which represents only modest levels. Using a lentiviral approach, we over-expressed VGLUT2 by only 50% (Fig 2), staying well within a physiological range within DA neurons. Consequently, we propose that the current strategy uniquely investigates the effect of slightly elevated VGLUT2 expression in DA neurons, as occurs after a lesion, thus promoting axon outgrowth. In contrast, in the previous report (Steinkellner et al., 2018), an AAV-infection was used, a strategy which is known to result in highly efficient gene expression (Saraiva et al., 2016). It is thus reasonable to assume that in that study, VGLUT2 over-expression levels were significantly above physiological levels and could therefore have initiated non-physiological effects leading to cell death (Steinkellner et al., 2018), perhaps resulting from abnormal intracellular phosphate levels. Indeed, one could speculate that adult DA neurons express only low levels (10-15 copies) of *Vglut2* mRNA to avoid toxicity, while simultaneously maintaining a sufficient baseline level to be able to reactivate this neurochemical phenotype when required (Fig 6A).

### Closing remarks

Previous work showed that an uptake inhibitor of the amino acid glycine, a co-agonist of the NMDA glutamate receptor, can promote reinnervation of the dorsal striatum after a 6-OHDA lesion (Schmitz et al., 2013). It was proposed that this mechanism could potentially be utilized in Parkinson’s disease patients to promote innervation of striatum by surviving DA neurons. The present work adds support to this possibility by demonstrating that Vglut2 expression in DA neurons and thus glutamate release by these neurons may represent an endogenous mechanism acting to facilitate reinnervation the striatum in parallel to the gradual loss of DA neurons during the course of this disease.

## Methods

### Animals

The experimental protocols were approved by the Animal Handling and Ethics Committee of the Université de Montréal (Montréal, QC, Canada). Housing was at a constant temperature (21^°^C) and humidity (60%), under a fixed 12h light/dark cycle and free access to food and water.

#### THGFP mice

Single cell qPCR experiments were performed using the tyrosine hydroxylase green fluorescent protein (THGFP) transgenic mouse line *TH-EGFP/21–31*, which carries the enhanced GFP (EGFP) gene under control of the TH promoter (Matsushita et al., 2002)

*Dat-Ires-Cre;AI9* Dat-Ires-Cre animals (Jax stock: 006660; (Bäckman et al., 2006)) were crossed with Ai9/tdTomato mice (Jax stock: 007905, a kind gift from prof. Williams at the Douglas Hospital).

#### Conditional Vglut2 knockout mice

6-OHDA and retrobead injection experiments were performed using conditional *Vglut2* knockout mice and control littermates. Dat-Ires-Cre animals (Jax stock: 006660) were crossed with Vglut2flox/flox mice (129, C57Bl/6 background) carrying the exon 2 surrounded by loxP site. A breeding colony was maintained by mating DAT-CRE;*Vglut2*^flox/+^ mice with *Vglut2*^flox/flox^ mice. 25% of the offspring from such mating were thus used as controls (i.e. DAT-CRE;*Vglut2*^flox/+^ mice) and 25% lacked *Vglut2* in DA neurons (i.e. DAT-CRE;*Vglut2*^flox/flox^ mice). P1-P2 mice were cryoanesthetized and decapitated for tissue collection. Primary cultures of mesencephalic DA neurons were prepared according to a previously described protocol (Fasano et al., 2008b). Mesencephalic cells were plated on monolayers of astrocytes at different densities, as described below.

#### Constitutive Vglut2 knockout mice

Mice heterologous for Vglut2 (maintained on a C57 background) were crossed to generate litters that included wild type, heterozygous and homozygous animals for the Vglut2 gene.

#### Intersectional genetic mice

Lineage fate-mapping experiments were performed using an intersectional strategy in which mice expressed Vglut2-Cre and TH-Flpo drivers in combination with AI65; as well as Vglut2-Cre and Dat-tTA in combination with AI82. More details on these mice and breeding strategies were published previously (Poulin et al., 2018).

### FACS-purified mesencephalic cultures

#### E11.5 cultures

THGFP-positive embryos were isolated at embryonic day E11.5, considering the morning of detection of the vaginal plug as E0.5. The mesencephalon was dissected and cells were prepared into a single cell suspension using a previously published protocol (Fasano et al., 2008). Cells were purified by fluorescence-activated cell sorting (FACS) using a BD FACSAria (BD BioSciences) and seeded in a 10k/ml concentration.

#### Postnatal cultures

The mesencephalon was dissected and cells taken from THGFP animals were dissociated to a single cell suspension using a previously published protocol (Fasano et al., 2008). Cells were purified by fluorescence-activated cell sorting (FACS). Cells were seeded at different concentrations for different experiments (table 1).

**Table 1.** In vitro treatment conditions. FACS: Before indicates that cells were sorted before seeding, while After indicates cells were sorted after the in vitro treatment.

| Name | Cell type | FACS | Concentration | Treatment | Incubation | Read-out |
| --- | --- | --- | --- | --- | --- | --- |
| Acute MPP+ | THGFP, mesencephalon | Before | 10.000/mL | 10 $\mu\text{M}$ | 24h | ICC |
| Chronic MPP+ | THGFP, SN | Before | 10.000/mL | 2 $\mu\text{M}$ | 72h | Single cell qPCR |
| Chronic MPP+ | Dat;Ai9, SN | After | 10.000/mL | 2 $\mu\text{M}$ | 72h | Bulk qPCR |
| GDNF | THGFP, mesencephalon | Before | 10.000/mL | 10 $\mu\text{M}$ | 3h | Single cell RT-PCR |

#### Vglut2-Venus over-expression

Vglut2 fused to Venus and Venus lentiviral vectors, using the Synapsin 1 promotor, were used for the over-expression of VGLUT2 in primary mesencephalic neurons. The vectors were a kind gift from Professor Etienne Herzog (University of Bordeaux), and were produced according to the strategy previously published (Herzog et al., 2011). Neurons were infected three hours after seeding (51316-106737 τυ/µl of Lenti-Venus (25% infected); 21000-24426 τυ/µl of Vglut2-Venus (42% infected)).

### Immunohistochemistry

#### Embryos

Coronal sections (12 μm) were cut on a cryostat and collected on Superfrost Plus microscope slides, after which they were subsequently washed with PBS, permeabilized, blocked and incubated overnight at 4°C with a rabbit anti-TH antibody (1:1000, AB152, Millipore Sigma, USA). Slices were washed the following morning and incubated for a minimum of 2h at room temperature (RT) with secondary antibody (1:1000, rabbit Alexa Fluor-546). Finally, sections were washed, counterstained with DAPI and were embedded with Fluoromount-G (Southern Biotech).

#### Neonates (P1 pups)

Coronal sections (25 μm) were cut on a cryostat were washed with PBS, permeabilized, blocked and incubated overnight at 4°C with primary anti-TH antibody (1:500, Pel-Freez P60101). Slices were washed the following morning and incubated for a minimum of 2h at RT with secondary antibody (rabbit Alexa Fluor-488 and rabbit Alexa Fluor-647, ThermoFisher). Finally, sections were washed, counterstained with DAPI and mounted on microscope slides.

#### Adult brain slices

Coronal sections (40 μm) were cut on a cryostat and stored in antifreeze solution at −20 °C until used. Slices were washed with PBS, permeabilized, blocked and incubated overnight at 4°C with rabbit anti-TH antibody (1:1000, AB152, Millipore Sigma, USA). Slices were washed the following morning and incubated for a minimum of 2h at RT with secondary antibody (1:1000, ant-rabbit Alexa Fluor-647, Invitrogen). Finally, sections were washed, counterstained with DAPI and mounted in Fluoromount-G (Southern Biotech) on Superfrost/Plus microscope slides.

### Immunocytochemistry on cell cultures

Cultures were fixed with 4% paraformaldehyde (PFA), permeabilized, and nonspecific binding sites were blocked. To study the (over-)expression of Vglut2 protein by post-natal DA neurons, a chicken anti-GFP (1:2000, GFP-1020; Aves Labs, Tigard, OR, USA), and a rabbit anti-VGLUT2 (1:2000, 135 402; Synaptic Systems, Göttingen, Germany). To study the expression of Vglut2 by embryonic DA neurons a mouse anti-VGLUT2 (1:4000, Millipore, AB5504) and primary anti-TH antibody (1:1000, AB152, Millipore Sigma, USA) were used. Cultures were washed the following morning and incubated for a minimum of 2h at RT with secondary antibodies (anti-rabbit Alexa Fluor-546/ anti-chicken Alexa Fluor-488 and rabbit Alexa Fluor-546/ anti-mouse Alexa Fluor-647/ anti-chicken Alexa Fluor-488 respectively, all 1:1000, Invitrogen). Finally, coverslips were washed, counterstained with DAPI and mounted in Fluoromount-G (Southern Biotech) on Superfrost/Plus microscope slides.

### Image acquisition with confocal microscopy

All *in vitro* images were acquired with a confocal microscope Olympus Fluoview FV1000 system (Olympus Canada, Markham, Ontario, Canada). For the quantification of Vglut2 protein in post-natal cells, 10 random field of views were chosen per coverslip and images were acquired using a 546 nm excitation laser and a 10x water immersion objective. For the analysis of the length of neuronal processes, images of 5 isolated, TH-positive neurons pre coverslip were acquired. For quantification of TH immunoreactivity in E18.5 dorsal and ventral striatum from Vglut2WT and Vglut2KO animals, two fields of view per hemisphere were acquired with a 60x oil-immersion objective. Three to four animals of each genotype were analyzed.

### Quantitative analysis of mesencephalon at E18.5

Images were taken with a Nikon Ti2 epifluorescence microscope, and using ImageJ software color combined images were split into single channel images, and subsequently converted into binary images (using default settings), after which the binary DAPI image was used as an overlay on top of the GFP image using image calculator. This creates a binary image of only the cells that were positive for both DAPI and GFP (cellular particles binary image), which were then counted using the ‘analyze particles’ function of ImageJ.

### Double fluorescent *in situ* hybridization (FISH)

THGFP-negative embryos were isolated at E11.5 and E14.5 and immediately frozen on dry ice. Sagittal slices of 10 µm were collected on SuperFrost Plus slides. Double FISH was performed as described previously (Fasano et al., 2017; Gras et al., 2008). Briefly, sections were fixed with 4% PFA (Tousimis) for 10 min, washed with PBS and acetylated with acetic anhydride for 10 min. Pre-hybridization was carried out by incubating the slides with hybridization mix (50% Formamide (Sigma), SSC (Ambion), E.coli tRNA (Roche), Denhardt’s, Salmon Sperm DNA) for 30 min at RT in a wet incubation chamber. Hybridization occurred over-night at 60 °C in a wet incubation chamber. After hybridization, slides were washed using several steps of SSC washes (2xSSC, 0.2xSSC) and incubated with blocking solution (FBS heat inactivated, blocking reagent (Roche). Probes were detected using a 1h incubation at RT with anti-fluorescein-POD (1:2500, Roche) and anti-DIG-POD antibodies (1:2500, Roche). Section were passed through several steps of washes in maleic acid buffer and in PBS-Tween (Fisher), after which the DIG-probes were visualized using TSA-Cy3 (PerkinElmer), and fluorescein-probes were visualized using TSA-Biotin (PerkinElmer) and Neutravidin-Oregon Green (Invitrogen). All sections were counterstained using DAPI and mounted using Fluoromount. Double FISH was performed with RNA probes labeled with digoxigenin (DIG) (Vglut2) and RNA probes labeled with Fluorescein (Th). Th probe was a kind gift from Prof. Marten Smidt (University of Amsterdam), and their sequences have been previously published (Grima et al., 1985; Jacobs et al., 2007). The sequence of the Vglut2 probe was published before (Herzog et al., 2001), and the probe was provided by from Prof. Salah El Mestikawy (McGill University, Montreal).

### Single-cell collection and qPCR

Experimental design was based on a previously described protocol that was published in great detail in (Fortin et al., 2019). In short, cultures were treated with 2.5 µM MPP+ at DIV11 and THGFP-positive cells were collected from coverslips using a glass pipette at DIV 14 and placed directly in an aliquot containing 0.5 ml of RNase Out (Thermo Fisher Scientific, Waltham, MA, USA) and 0,5 ml of DTT (Thermo Fisher Scientific). Then, total RNA from each cell was reverse transcribed in a total of 20 µl cDNA. Quantitative PCR was performed using SYBER Green PCR Master Mix (Quantabio, Beverly, MA, USA), on a Light Cycler 96 machine (F. Hoffmann-La Roche, Basel, Switzerland). The presence of a cell in each sample was confirmed by detecting the presence of glyceraldehyde-3-phosphate dehydrogenase (GAPDH) mRNA. Calculation of absolute copy number for TH and VGluT2 in each cell was determined with an external calibration curve based on a TH or VGluT2 plasmid cDNA (obtained, respectively, from Sino Biological (Beijing, China) and Harvard University (Boston, MA, USA). Primers were synthesized by AlphaDNA. Primers for qPCR were: TH 5’-TGGCCTTCCGTGTGTTT-3’ and 5’-AATGTCCTGGGAGAACTGG-3’; Vglut2 5’-CCTTTTGTGGTTCCTATGCT-3’ and 5’-GCTCTCTCCAATGCTCTCC-3’; GAPDH 5’-GGAGGAAACCTGCCAAGTATGA-3’ and 5’-TGAAGTCGCAGGAGACAACC-3.

### Embryo collection and tissue preparation

Embryos from several mouse lines were isolated at embryonic day (E)11.5, E14.5, and E18.5, considering the morning of detection of the vaginal plug as E0.5. Embryos were isolated in ice cold PBS and fixed in 4% PFA for 24-48h for immunohistochemistry, or freshly frozen on dry ice for in situ hybridization. Vglut2WT and Vglut2KO embryos were cryoprotected and shipped from the laboratory in Berlin (Germany) to Montreal (Canada) in 30% sucrose in PBS, on room temperature. Upon arrival embryos were stored immediately on 4°C and frozen on dry ice before cryostat sectioning (12 µm).

### Surgeries and tissue preparation

The 6-OHDA surgery protocol used here was previously described (Giguère et al., 2019). In short, DatICre;Vglut2WT and DatICre;Vglut2KO animals (2-6 months old) were anesthetized with isoflurane (Aerrane; Baxter, Deerfield, IL, USA) and fixed on a stereotaxic frame (Stoelting, Wood Dale, IL, USA). Animals received 6-OHDA injection (0.5 μL of 6-OHDA (3 mg/mL) in 0.2% ascorbic acid solution) in the dorsal striatum (Bregma coordinates: AP 0.5 mm; ML 2.0 mm; DV −3.0 mm). The norepinephrine transporter blocker, desipramine (2.5mg/ml; 10 µl per gram), was injected 45 min prior to 6-OHDA. Animals were allowed to recover in their own cages. After 6 weeks, animals received 100nl red fluorescent retrobeads IX (Lumafluor, Inc.) (1:1 diluted in saline) using the same surgery and anaesthesia protocol and bregma coordinates. One week after the retrobread injections, animals were anesthetized using pentobarbital NaCl saline solution (7 mg/mL) injected intraperitoneally, after which they were perfused with 50mlL of PBS followed by 100 ml of PFA 4% using an intracardiac needle at a rate of 25 ml/min. Brains were post-fixed in 4% PFA for 48h, transferred to 30% sucrose in PBS solution and stored at −80 °C before being sectioned at the cryostat at 40 µm.

### Analysis of Rb+/TH+ cells in mesencephalon

The striatum was imaged using the Nikon Ti2 microscope using a 20x objective. Using Nikon Elements software, the circumference of Rb-fluorescence at the injection site was traced to calculate the area. The number of counted Rb-positive cells in the mesencephalon of each mouse was divided by the area of Rb in the striatum to normalize for variation in the injection volume. Stacks of 5um were acquired (5-6 slices per stack), taking a 2mm by 2mm field of view of the ipsilateral injection side, for TH (Alexa 647) and retrobeads (RFP). Stitched images were analyzed as maximum intensity projections. TH-negative and TH-positive cells containing Retrobeads were counted blindly, manually.

### Analysis of TH immunoreactivity (IR) in dorsal striatal sections

#### Fluorescence TH-IR

The striatal slices were imaged using the Nikon Ti2 microscope using a 5x objective. Using Nikon Elements software, the circumference of the dorsal striatum were traced, in both ipsi- and contra-lateral side and determine the average TH-IR in each region of interest. The average TH-IR of each ipsilateral site was divided by the average TH-IR of the contralateral side. These values were used to obtain the average TH-IR ratio per brain, to quantify the lesion size per brain. Lesions were considered successful if TH-IR was reduced by two times the standard deviation of saline control values (i.e. 7%).

#### DAB TH-IR

The striatal slices were imaged using the Micromanager Nikon Eclipse TE200 inverted microscope with 10x objective (N.A. 0.25) and analyzed by a blinded observer. ImageJ was used to draw the circumference of the dorsal striatum were traced, in both ipsi- and contra-lateral side and to determine the average TH-IR in each region of interest. The average TH-IR of each ipsilateral site was divided by the average TH-IR of the contralateral side. These values were added to obtain the total TH-IR ratio per brain, to quantify the lesion size per brain.

### Stereology and DAB immunohistochemistry

Brain slices (40 µm) of Vglut2cWT and Vglut2cKO animals that had undergone surgery were washed with PBS and the endogenous peroxidase activity was quenched with 0.9% hydrogen peroxidase. Slices were permeabilized and non-specific binding sites were blocked before incubation with rabbit anti-TH antibody (1:1000, AB152, Millipore Sigma, USA) for 48h. Subsequently, slices were incubated at 4°C for 12h with goat anti-rabbit biotin-SP-AffiniPure secondary antibody (111-065-003, 1:200, Jackson ImmunoResearch Laboratories, USA), and finally for 3h with horseradish peroxidase streptavidin (016-030-084, Cedarlane, USA). The DAB reaction was carried out for 15s, then stopped by incubation with 0.1M acetate buffer and slices were mounted on Superfrost/Plus microscope slides. They were left to dry for 96h after which they were counterstained with cresyl violet, dehydrated and finally slides were sealed with Permount mounting medium (SP15-100, Fisher, USA) using glass coverslips.

### Stereological counting

SN TH-immunoreactive neurons were counted in every sixth section using a 100x oil-immersion objective on a Leica DMRE stereomicroscope equipped with a motorized stage by blinded observer. A 60 x 60 μm^2^ counting frame was used in the Stereo Investigator (MBF Bioscience) sampling software with a 20 μm optical dissector (2 μm guard zones) and counting site intervals of 100 μm after a random start. Stereological estimates of the total number of TH-immunoreactive neurons within ipsi-lateral SN were obtained, and these numbers are presented relative to the number of TH-positive neurons counted on the intact contra-lateral side.

### Statistics

Data are represented throughout as means ± SEM. Statistically significant differences were analyzed with Student’s t test, 2-way ANOVA (Tukey’s post-hoc), or a Fisher’s exact test, as appropriate. P<0.05 was considered significant (*), P<0.01 (**), P<0.001 (***).

## Acknowledgements

We thank Dr. Alexander Weil for access to the StereoInvestigator stereological analysis workstation, Dr. Daniel Lévesque for access to his laboratory space and cryostat and Dr. Sebastian Talbot for access to his Nikon Ti2 epifluorescence microscope. We thank Dominic Thibault for his technical assistance with the viral preparations.

